# Inference for entomological semi-field experiments: Fitting a mathematical model assessing personal and community protection of vector-control interventions

**DOI:** 10.1101/2023.06.30.547223

**Authors:** Emma L Fairbanks, Manop Saeung, Arissara Pongsiri, Elodie Vajda, Yuqian Wang, David J McIver, Allison Tatarsky, Neil F Lobo, Sarah J Moore, Alongkot Ponlawat, Theeraphap Chareonviriyaphap, Amanda Ross, Nakul Chitnis

**Affiliations:** Department of Epidemiology and Public Health, Swiss Tropical and Public Health, Institute, Allschwill, Switzerland; University of Basel, Basel, Switzerland; Department of Entomology, Faculty of Agriculture, Kasetsart University, Bangkok, Thailand; Department of Entomology, Armed Forces Research Institute of Medical Sciences (AFRIMS), Bangkok, Thailand; Malaria Elimination Initiative, Institute for Global Health Sciences, University of California, San Francisco; University of Notre Dame, Indiana, USA; Vector Control Product Testing Unit, Ifakara Health Institute, Bagamoyo, United Republic of Tanzania; The Nelson Mandela, African Institution of Science and Technology, School of Life Sciences and Bio Engineering, Tengeru, Arusha, United Republic of Tanzania

**Author notes:** Corresponding author; Swiss Tropical and Public Health Institute, Kreuzstrasse 2, 4123 Allschwil, Switzerland.

**Keywords:** Malaria, *Anopheles*, vector control, spatial repellent, volatile pyrethroid, picaridin personal protection, community protection, insecticide-treated clothing

## Abstract

The effectiveness of vector-control tools is often assessed by experiments as a reduction in mosquito landings using human landing catches (HLCs). However, HLCs alone only quantify a single characteristic and therefore do not provide information on the overall impacts of the intervention product. Using data from a recent semi-field study which used time-stratified HLCs, aspiration of non-landing mosquitoes, and blood feeding, we suggest a Bayesian inference approach for fitting such data to a stochastic model. This model considers both personal protection, through a reduction in biting, and community protection, from mosquito mortality and disarming (prolonged inhibition of blood feeding). Parameter estimates are then used to predict the reduction of vectorial capacity induced by etofenpox-treated clothing, picaridin topical repellents, transfluthrin spatial repellents and metofluthrin spatial repellents, as well as combined interventions for *Plasmodium falciparum* malaria in *Anopleles minimus*. Overall, all interventions had both personal and community effects, preventing biting and killing or disarming mosquitoes. This led to large estimated reductions in the vectorial capacity, with substantial impact even at low coverage. As the interventions aged, fewer mosquitoes were killed; however the impact of some interventions changed from killing to disarming mosquitoes. Overall, this inference method allows for additional modes of action, rather than just reduction in biting, to be parameterised and highlights the tools assessed as promising malaria interventions.

## 1 Introduction

Almost half of the population at risk of malaria lives in forested areas [17]. These forests are ideal conditions for many vectors, due to the vegetation cover, temperature, rainfall and humidity [25]. Tools such as indoor residual spraying (IRS) and long-lasting insectcidal nets (LLINs) are not effective against outdoor biting, which is associated with occupations within forested regions [2, 28]. Volatile pyrethroid spatial repellents and insecticide-treated clothing are promising new tools for bite prevention in these regions [1].

Permethrin-treated military uniforms (PTMUs) have been shown to reduce biting of *Anopheles dirus* and *Anopheles farauti* in laboratory bioassays [12, 14] and experimental huts in Cote d’Ivoire [11]. However, knockdown (incapacitation) and mortality reduced significantly in laboratory settings when PTMUs were washed three times for *Anopheles dirus* [12] and *Anopheles farauti* [13], although this is not observed for all pyrethroid-treated fabrics [13].

In epidemiological trails PTMUs did not offer enough protection to significantly reduce malaria incidences in troops with low-or-no immunity the high-incidence settings of Cote d’Ivoire [11] or northeastern Thailand [12]. However, little is known about the potential effects of these interventions in low-incidence settings or in populations with immunity. There are limited studies on repellent-treated clothing or treated civilian clothing, however transfluthrin-treated sandals have been shown to reduce *Anopheles arabiensis* bites by 54-86% [22]. In Cote d’Ivoire topical repellent (50% DEET) combined with the PTMUs did not significantly reduce malaria incidences [11]. However, in laboratory conditions DEET-treated mosquito netting has been shown to repel, inhibit blood-feeding and kill mosquitoes for at least 6 months [23].

Modelling studies parameterising the impact of vector-control tools have typically relied on data from experimental-hut studies [4, 29], as well as experimental-hut data combined with laboratory bioassay data [8]. If available, these models can be validated by comparing simulated results to randomised control trial data [29]. Mathematical models have also been fit to entomological field trial data to assess the potential impact of the novel intervention Attractive Targeted Sugar Baits to reduce malaria transmission [15, 19]. These models generally try to describe how the vector-control tools influence the mosquitoes feeding cycles (Figure 1).

**Figure 1.**
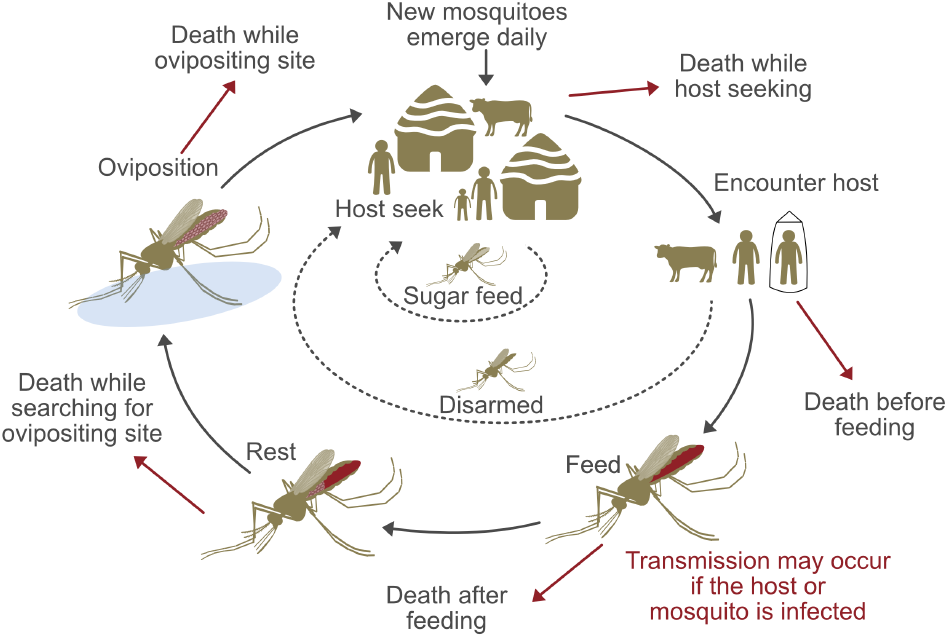
: The *Anopheles* feeding cycle. Mosquitoes emerge at the start of the cycle, the host-seeking stage. From here they will either die, sugar feed or encounter a host. If they encounter a host they may die before feeding, be disarmed (inhibited from blood feeding) or feed. If they feed malaria transmission may occur to the host or mosquito. After feeding the mosquito dies or rests while their eggs develop. Then, they will search for a site for oviposition. If they survive this search they will lay the eggs and either die or return to the host-seeking stage and begin the cycle again. Figure modified from [7].

Recently, semi-field sites have been developed that allow us to determine the impacts of tools on entomological outcomes in a controlled environment [10, 20, 31, 38]. These studies involve releasing a known population of a mosquitoes into a cage and typically use a standard measure of human landing catches (HLCs) to assess the potential impact of interventions, measuring the change in mosquitoes landing on a human [36]. The HLC method allows insight into the personal protection of the tools for the user, however it does not offer any information as to why there is a change in the rate of landing. There are many possible ‘end points’ for these mosquitoes. Considering these additional end points allows assessment of potential community protection, which is essential to understanding the full impact of an intervention [18]. Here, using semi-field study data, we further investigate the ‘modes of action’ of vector-control tools which lead to this community protection. We consider preprandial mortality, disarming, postprandial mortality, and repellency of mosquitoes. If an infectious mosquito is killed preprandially (before biting), it cannot transmit malaria to a host. If a mosquito is disarmed it is inhibited from blood feeding and therefore cannot transmit malaria during the current feeding cycle, however, if infected, it may continue the extrinsic incubation period and infect humans in future feeding cycles. An infectious mosquito that dies postprandially (after biting) may transmit malaria to the host on which it fed. If a mosquito is repelled it remains in the host seeking stage and may bite another host during the current feeding cycle. There will be community protection from preprandial and postprandial killing and disarming, since these end points prevent the mosquito biting other members of the community during the current (and future if the mosquito is killed) feeding cycle, unlike repellency.

Denz et al. [10] developed models and parameterisation methods to estimate the effect of spatial repellents and odour-baited traps on the rate at which mosquitoes fed on humans based on only HLC data from a semi-field study [24]. Although model allowed differentiation between repellency, preprandial mortality and disarming, data allowing for parameterisation of the magnitude of these effects was not available. It was assumed that all the reduction in HLCs was due to preprandial killing or disarming. the study also modelled and parameterised postprandial killing based on the proportion of mosquitoes caught by HLC which died within 24 hours. Results from these models were then used to inform a previously published model for vectorial capacity [7], estimating the potential impact of these interventions. Their results showed that the distinction disarming and preprandial mortality can have large impacts on estimates of reduction in vectorial capacity.

In this study, we use data from semi-field experiments extended to include aspiration of uncaught resting mosquitoes after the HLC collection period, and the offering of blood meals to all HLC caught and resting mosquitoes to determine subsequent feeding probability. A novel statistical method, considering these additional data, is suggested that, for the first time, allows us tor parameterise the change in host-availability, disarming and preprandial mortality rates. Estimation for the decrease in biting, increase in postprandial and preprandial mortality and disarming effects of interventions on mosquitoes are used to provide more refined predictions of the relative reduction in vectorial capacity for a range of coverage levels within a population for each intervention.

## 2 Semi-field study data

Semi-field experiments were carried out at the Pu Teuy mosquito field research station (14°20N; 98°59E), Kasetsart University (KU) and The Armed Forces Research Institute of Medical Sciences (AFRIMS), Kamphaeng Phet. The experiments were performed for 8 nights per intervention arm, per site, with control and intervention experiments ran in parallel each night. Initial experiments were carried out July 2020–Janurary 2021. This was followed by further experiments in May–July 2022. A summary of the data collected each experiment is given in Table 1.

**Table 1:**
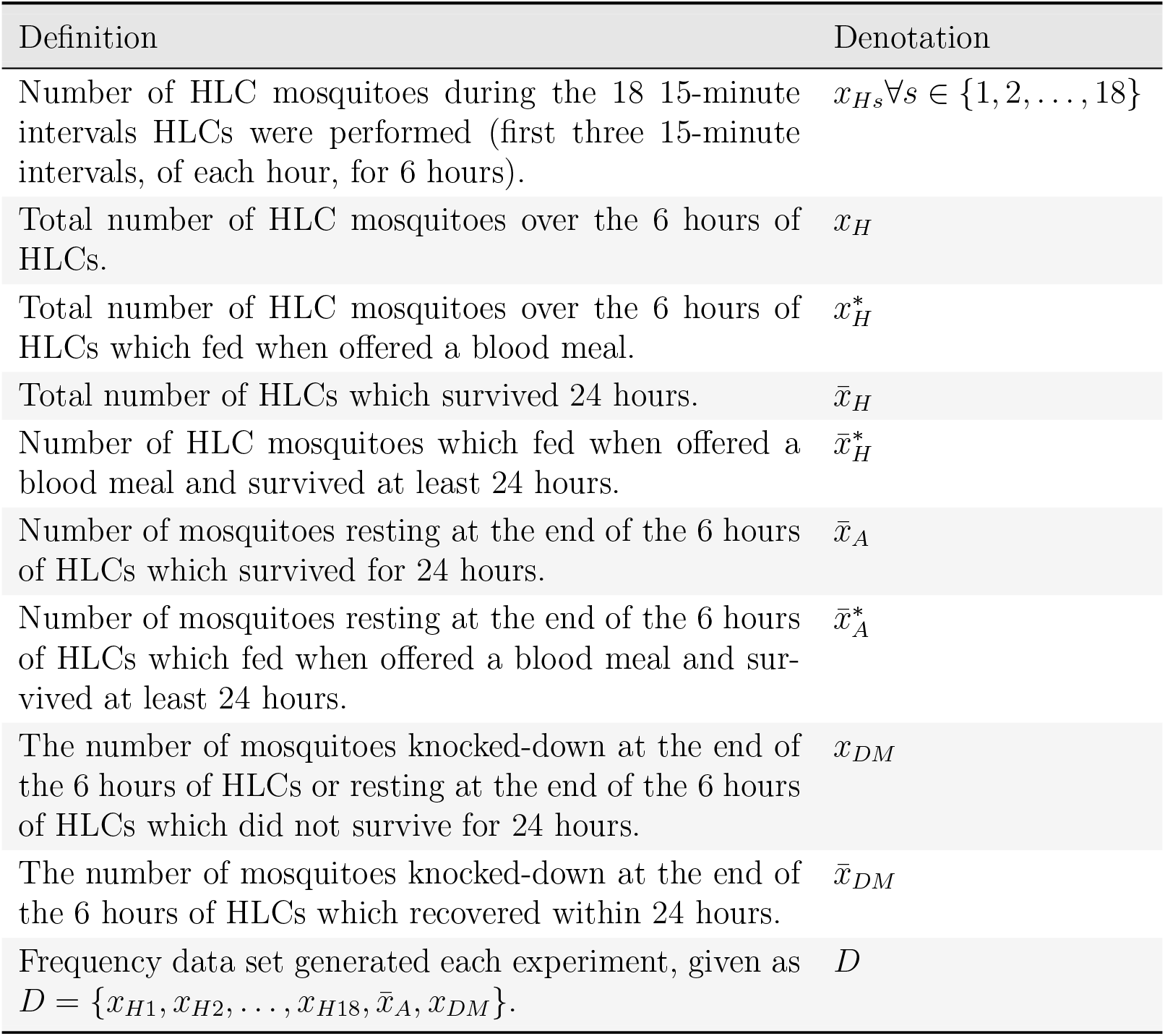
Data generated from semi-field studies used in the statistical inference.

### 2.1 Interventions

Nine combinations of transfluthrin spatial repellents, metofluthrin spatial repellents, clothing treated with the pyrethroid insecticide etofenprox and the aerosol, spray-on topical repellent picaridin (20%) were considered:

1. Etofenprox-treated civilian clothing — shorts and short-sleeved t-shirt (with bare lower arms and legs), referred to as EtoCivilianA.
2. Etofenprox-treated civilian clothing with picaridin, referred to as EtoCivilianAPi.
3. Etofenprox-treated civilian clothing with picaridin — cotton with long trousers and short-sleeved t-shirt, referred to as EtoCivilianBPi.
4. Etofenprox-treated ranger clothing — full leg/arm length, referred to as EtoRanger.
5. Etofenprox-treated ranger clothing with picaridin, referred to as EtoRangerPi.
6. Passive transfluthrin spatial repellent — a hung treated piece of fabric, referred to as TransPassive.
7. Active (aerosol) transfluthrin spatial repellent, referred to as TransActive.
8. Passive metofluthrin spatial repellent — a hung treated piece of fabric, referred to as MetoPassive.
9. Etofenprox-treated civilian clothing with picaridin and passive transfluthrin spatial repellent, referred to as EtoCivilianBPi + TransPassive.

During all experiments volunteers wore closed shoes or boots. For treated clothing interventions there were two intervention arms: new and washed-20-times. This is denoted with an appended 0 or 20, for new and aged clothing, respectively. TransPassive was also trialed both new and aged-30-days, denoted with an appended 0 or 30, respectively. Table S1 shows which sites and years when experiments on each intervention were performed.

### 2.2 Experiment protocol

100 insectory-reared, aged-5–8-days *Anopheles minimus s*.*s*. which had never been offered a blood meal and had been starved of sugar for 8 hours prior to the experiment, were released into screen houses. Volunteers performed HLCs in the screen houses for 6 hours. Mosquitoes caught by HLC were placed in a collection cup. The number of mosquitoes collected by HLCs was recorded every 15 minutes for the first 45 minutes of each hour; then during the final 15 minutes, the three collection cups for that hour were removed and the collectors rested outside the screen house. This was to avoid contamination of the mosquitoes due to close proximity to the interventions while in the collection cup, and also improved the comfort and alertness of volunteers. At the end of the 6-hour landing period the remaining mosquitoes were collected by Prokopack aspiration. The number of knocked-down and resting mosquitoes collected by aspiration was recorded. HLC and resting mosquitoes were subsequently offered a blood meal for 30 minutes using the membrane technique. This was performed in cups containing a maximum of 20 mosquitoes. After the blood meal was offered, each mosquito cup received 10% sucrose soaked cotton balls. Mosquitoes were examined after 24 hours and whether they were fed and alive, fed and dead, unfed and alive or unfed and dead was recorded. Knocked-down mosquitoes were also examined after 24 hours. Here, whether they remained knocked-down and were therefore classified as dead, or recovered and were therefore classified as disarmed was recorded [35].

Mosquitoes knocked-down or resting at the end of the 6-hours of HLCs which were dead after 24 hours were classified as dead. It is assumed that these mosquitoes would not have survived until the next days feeding period, and therefore would have died before feeding on a human and potentially transmitting malaria.

During round one of the experiments (2020–2021) relatively low levels of blood feeding were recorded during many of the control experiments. The World Health Organisation (WHO) guidelines for efficacy testing of spatial repellents suggest that 50% of control mosquitoes should land and 25% should feed [39]. These behaviours are desired in the control as a change in these behaviours is needed to measure the effect of the active ingredient. A lack of these behaviours may be due to an error in the rearing process or damage by HLC volunteers during capture. WHO guidelines on ITN evaluation suggest 50% of control mosquitoes should feed [40]. This is because mosquitoes that blood feed are more likely to survive. Since we also want to measure postprandial mortality here, we have combined these guidelines and introduced a 50% threshold for landing and feeding. This was included in the Standard Operating Procedure in round two of the experiments. For consistency, we only consider data which meets this criteria for both rounds of this study. Table S1 describes how many experiments met this specification for each intervention and year.

## 3 Methods

Figure 2 shows a schematic of the processes being modelled. Section 3.1 defines how each process is modelled, with Table 2 defining all parameters in the section. Section 3.2 then describes how these models are given a hierarchical Bayesian structure allowing for variation in behaviour between experiments.

**Table 2:** Parameters for the mathematical and statistical models described in Section 3.1. *k* denotes the night for parameters that vary nightly. HLC = human landing catches.

| Parameter | Definition |
| --- | --- |
| $t$ | time-step (with units of one hour) |
| $P_A(t)$ | Probability mosquito remains in the host-seeking stage during time-step |
| $P_H(t)$ | Probability mosquito is caught by HLC during time-step |
| $P_{DM}(t)$ | Probability mosquito is killed or disarmed during time-step |
| $\alpha_H$ | Rate host-seeking mosquitoes are caught by HLC ($\text{hour}^{-1}$) |
| $\alpha_{DM}$ | Rate host-seeking mosquitoes killed or disarmed ($\text{hour}^{-1}$) |
| $P_{F,k}$ | Probability a mosquito feeds night $k$ |
| $P_{FA,k}$ | Probability a resting mosquito feeds night $k$ |
| $P_{FH,k}$ | Probability a HLC mosquito feeds night $k$ |
| $P_{D,k}$ | Probability a mosquito from the intervention experiment is disarmed on night $k$ |
| $P_{DA,k}$ | Probability a resting mosquito from the intervention experiment is disarmed on night $k$ |
| $P_{DH,k}$ | Probability a HLC mosquito from the intervention experiment is disarmed on night $k$ |
| $\pi$ | Relative reduction in the host-availability rate for mosquitoes encountering a human protected by the intervention compared to a unprotected (control) human |
| $\kappa$ | Increase in the rate of preprandial mortality or disarming due to an intervention relative to the rate of HLCs in the control |
| $\xi$ | Increased probability of death after a mosquito has fed on a human protected by the intervention |
| $P_{M,k}$ | Probability of mortality after a mosquito bites a human |
| $\omega$ | Proportion of mosquitoes disarmed of all mosquitoes either killed preprandially or disarmed |

**Figure 2.**
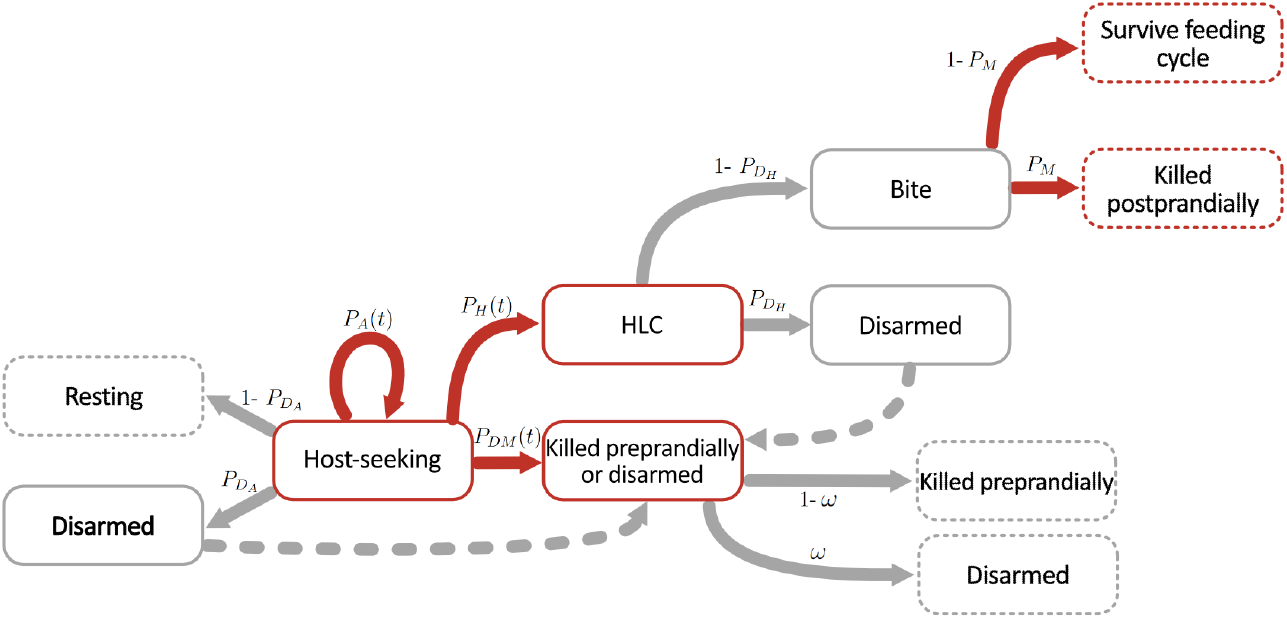
: Schematic of the processes modelled. Red boxes and arrows include processes previously modelled and parameterised in Denz et al. [10]. Grey boxes and arrows show extensions novel to this work. Definitions of the probabilities associated with each process are given in Table 2. Boxes with a dashed outline represent a possible end-point for mosquitoes. Dashed lines represent were resting and HLC mosquitoes are reclassified as disarmed based off blood-feeding data.

Within each intervention arm we denote nightly values of parameters and data points with the subscript *k*, where *k* ∈ {1, 2, …, 8}. Parameters and data points referring to the control and intervention experiments are denoted [*C*] and [*I*], respectively.

Figures were generated using the *ggplot2* package in *Rstudio* [27, 37].

### 3.1 Mathematical and statistical models

#### 3.1.1 Modelling the individual behaviour of mosquitoes encountering unprotected humans

We consider a stochastic continuous-time Markov model of host-seeking behaviour at the individual mosquito level [7]. The mosquito starts in the host-seeking stage (*A*). During a given time step (duration *t*) a mosquito can either remain in *A*, be caught by HLC (*H*) or be killed or disarmed (*DM*), with probabilities *P*_*A*_(*t*), *P*_*H*_ (*t*) and *P*_*DM*_ (*t*), respectively. It is assumed that these probabilities are independent of how long the mosquito has been in state *A* and mosquitoes leave state *A* following an exponential distribution. We assume that the probabilities that a mosquito moves to state *H* or *DM* within a short time can be approximated linearly in time with constant rates *α*_*H*_ and *α*_*DM*_, respectively. Here we consider the time step duration (*t*) to be one hour.

Each time step the probability that a mosquito remains in state *A* is

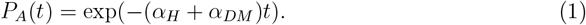

Thus, the probability that a mosquito leaves state *A* is given as 1 −*P*_*A*_(*t*). Given a mosquito left state *A* it then enters state *H* or *DM* with probabilities

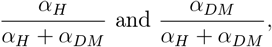

respectively. Therefore, the probabilities that a mosquito moves to state *H* and *DM* are

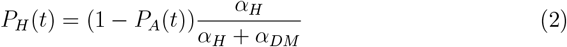

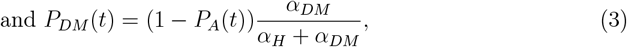

respectively.

#### 3.1.2 Modelling the differences in individual behaviour of mosquitoes encountering protected and unprotected humans

##### Statistical model for the probability a mosquito is disarmed

Since we assume there is no disarming in the control experiment, the probability a control mosquito feeds on night *k* (*P*_*F,k*_[*C*]) is equivalent to the probability that a non-disarmed mosquito feeds. The probability a mosquito feeds in the intervention experiment *P*_*F,k*_[*I*] is defined as the probability the mosquito is not disarmed × the probability a non-disarmed mosquito feeds. Therefore, *P*_*F,k*_[*I*] = (1 −*P*_*D,k*_)*P*_*F,k*_[*C*], where *P*_*D,k*_ describes the probability a mosquito from the intervention experiment, offered a blood meal on night *k* is disarmed. This proportion may not be equal for resting and HLC mosquitoes. We can allow for different feeding and disarming probabilities for resting and HLC mosquitoes, denoted 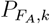 and 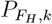 and 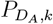 and 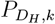 respectively (Table 2).

##### Change in host-availability rate

We define a parameter *π* ∈ [0, 1] which describes the relative reduction in the host-availability rate for mosquitoes encountering a human protected by the intervention compared to an unprotected (control) human. Then, the nightly host availability rate for the protected host is

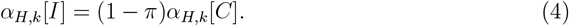

##### Change in preprandial mortality or disarming rate

In addition to reducing HLCs interventions may also increase the rate of preprandial mortality or disarming. We define *κ* ∈ [0, ∞) as the increase in the rate of preprandial mortality or disarming due to an intervention relative to the host-availability rate in the control. Then, the nightly rate of intervention preprandial mortality or disarming is

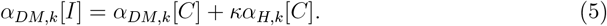

##### Statistical model for postprandial mortality

The postprandial mortality effect, *ξ* ∈ [0, 1], is defined as the increased probability of death after a mosquito has fed on a human protected by an intervention. We parameterise the postprandial mortality affect using the model suggested by Denz et al. [10]. Briefly, the probability of mortality after a mosquito bites a protected human on day *k* is given by

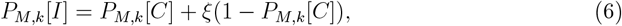

where *P*_*M,k*_[*C*] is the probability of mortality after a mosquito bites an unprotected human.

##### Contribution of preprandial mortality vs. disarming towards *κ*

The contributions to *κ* from preprandial mortality and disarming can be estimated according to the ratio of disarmed to mosquitoes killed preprandially. We introduce a new parameter *ω* defined as

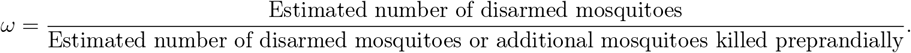

The contribution to kappa from disarming and additional preprandial killing are then estimated as *ω × κ* and (1 − *ω*) × *κ*, respectively.

To calculate the increase in mosquitoes killed preprandially we consider:

i. The increase in the number of mosquitoes knocked-down after the 6 hours of HLCs which do not recover after 24 hours or mosquitoes resting at the end of the 6 hours of HLCs which did not survive for 24 hours, from the control to intervention arms 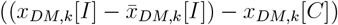.
ii. The disarmed HLC mosquitoes which die within 24 hours. The number of HLCs which do not survive 24 hours is calculated as 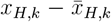 these are a combination of mosquitoes killed postprandially and mosquitoes disarmed when they were offered a blood meal which do not survive 24 hours. The number of HLCs that are not disarmed in this group can be estimated as number of HLCs which are not disarmed and killed postprandially, given as 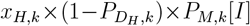. Therefore, the number of mosquitoes which were disarmed at the time of blood feeding but do not survive to the next feeding cycle, therefore, classed as preprandially killed mosquitoes, is calculated as 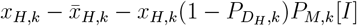.

To calculate the increase in mosquitoes disarmed we consider:

i. The number of disarmed resting mosquitoes, estimated as 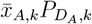.
ii. The number of mosquitoes knocked-down after the 6 hours of HLC which recover after 24 hours 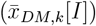.
iii. The disarmed HLC mosquitoes which do not die within 24 hours. This is estimated as the number of disarmed HLCs 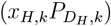 without the disarmed HLC mosquitoes which die (estimated above), altogether giving 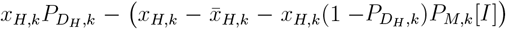.

### 3.2 Hierarchical structure allowing for variation in behaviour between experiments

In order to consider the nightly variations in the parameters we consider a Bayesian hierarchical model.

#### 3.2.1 Host-availability and preprandial mortality or disarming rates

Applying a hierarchical Bayesian model we allow *α*_*H*_ and *α*_*DM*_ to vary between nights. The nightly rates for the control experiments are assumed to follow the distributions

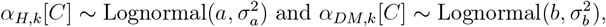

where *a, b* ∈ ℝ and *σ*_*a*_, *σ*_*b*_ ∈ ℝ^+^ are defined in Table 3. The mean rates are the expected value of these distributions, calculated as

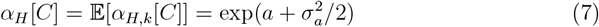

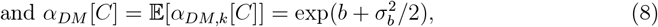

respectively.

**Table 3:** Hyperparamters in the Bayesian hierarchial model. *k* denotes the night for parameters that vary nightly.

| Parameter | Definition | Prior |
| --- | --- | --- |
| Probabilities of feeding and disarming for HLC and resting mosquitoes |  |  |
| $u$ | $\text{logit}(P_{F,k}[C])$ mean | Logistic(0,1) |
| $\sigma_u$ | $\text{logit}(P_{F,k}[C])$ standard deviation | Half-Cauchy(0,1) |
| $\phi_{u_k}$ | Normalised deviation of $\text{logit}(P_{F,k}[C])$ from mean | $\mathcal{N}(0, 1)$ |
| $v$ | $\text{logit}(P_{D_A,k}[I])$ mean | Logistic(0,1) |
| $\sigma_v$ | $\text{logit}(P_{D_A,k}[I])$ standard deviation | Half-Cauchy(0,1) |
| $\phi_{v,k}$ | Normalised deviation of $\text{logit}(P_{D_A,k}[I])$ from mean | $\mathcal{N}(0, 1)$ |
| $w$ | $\text{logit}(P_{D_H,k}[I])$ mean | Logistic(0,1) |
| $\sigma_w$ | $\text{logit}(P_{D_H,k}[I])$ standard deviation | Half-Cauchy(0,1) |
| $\phi_{w,k}$ | Normalised deviation of $\text{logit}(P_{D_H,k}[I])$ from mean | $\mathcal{N}(0, 1)$ |
| Host-availability and preprandial mortality or disarming rates |  |  |
| $a$ | $\log(\alpha_{H,k})$ mean | $\mathcal{N}(0, 6)$ |
| $\sigma_a$ | $\log(\alpha_{H,k})$ standard deviation | Half-Cauchy(0,1) |
| $\phi_{a,k}$ | Normalised deviation of $\log(\alpha_{H,K})$ from mean | $\mathcal{N}(0, 1)$ |
| $b$ | $\log(\alpha_{DM,k})$ mean | $\mathcal{N}(0, 6)$ |
| $\sigma_b$ | $\log(\alpha_{DM,k})$ standard deviation | Half-Cauchy(0,1) |
| $\phi_{b,k}$ | Normalised deviation of $\log(\alpha_{DM,K})$ from mean | $\mathcal{N}(0, 1)$ |
| $1 - \pi$ | Intervention effect on host-availability rate | Lognormal(0,5) |
| $\kappa$ | Intervention effect on preprandial mortality or disarming rate | Lognormal(0,5) |
| Probability of postprandial mortality |  |  |
| $r$ | $\text{logit}(P_{M,k}[C])$ mean | Logistic(0,1) |
| $\sigma_r$ | $\text{logit}(P_{M,k})$ standard deviation | Half-Cauchy(0,1) |
| $\phi_{r_k}$ | Normalised deviation of $\text{logit}(P_{M,k})$ from mean | $\mathcal{N}(0, 1)$ |
| $\xi$ | Intervention effect on postprandial mortality | Uniform(0,1) |

We consider *ϕ*_*a*_ and *ϕ*_*b*_, with elements *ϕ*_*a,k*_, *ϕ*_*b,k*_ ∈ ℝ, describing how *a* and *b* deviate nightly from the mean according to the scale of the standard deviation, that is

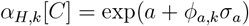

and

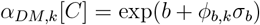

#### 3.2.2 Statistical model probabilities

The mean probabilities of a non-disarmed mosquito feeding (*P*_*F*_ [*C*]), a HLC mosquito is disarmed 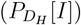, a resting mosquito is disarmed 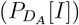 and a mosquito encountering an unprotected host dies after biting (*P*_*M*_ [*C*]) follow a logistic distribution. For a given logistically distributed parameter, *Z*, the nightly probability of the event is given as

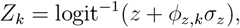

where 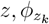 and *σ*_*z*_ ∈ R^+^ are hyperparameters. Specifically, *z* is the mean of logit(*Z*_*k*_), *σ*_*z*_ is the standard deviation of logit(*Z*_*k*_) and *ϕ*_*z,k*_ is the normalized deviation of logit(*Z*_*k*_) from the mean on night *k*.

The mean probability over all nights is calculated as

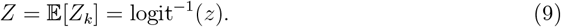

### 3.3 Bayesian Inference

Bayesian inference was performed in Stan [30] in Rstudio [27], which was used to generate figures. Weakly informed priors, similar to [10], were used (Table 3). For each model parameterised we run four Markov chains with 10,000 iterations, removing the first 5000 for burn-in. The convergence of chains was checked using the diagnostics available within Stan, including the R-statistic and effective sample size. Hyperparameter posteriors were plotted as pairwise scatter plots to identify any potential parameter identifiabiliy issues.

#### 3.3.1 Probabilities HLC and resting mosquitoes are disarmed

Given the number of mosquitoes in 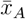 and (*x*_*H*1_, *x*_*H*2_, …, *x*_*H*18_) which fed when offered a blood meal are denoted 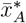 and 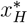, respectively, we denote the nightly values for the number of resting and HLC mosquitoes offered a blood meal 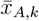 and *x*_*H,k*_, respectively, and the corresponding number which fed 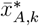 and 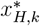, respectively. We then have

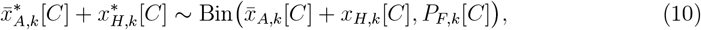

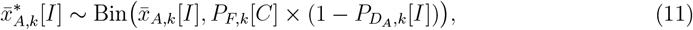

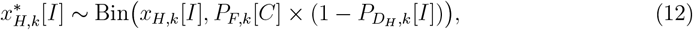

where 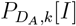 and 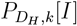 describe the probability a resting and HLC mosquito is disarmed, respectively. Here, in Equation 10, we assume that in the absence of interventions (control arm), the probability of feeding for resting and HLC mosquitoes is the same. This is because many control experiments had low counts of resting mosquitoes.

#### 3.3.2 Host-availability and preprandial mortality or disarming rates

We denote the 15-minute intervals in which HLC data is collected as 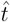, with 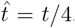. The probability that a mosquito remains in state *A* for the first *i* 15-minute intervals, and then enters state *H* during the following *j* 15-minute intervals is given by 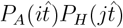. Since exp(*nx*) = exp(*x*)^*n*^, where *n* ∈ ℤ^+^, it follows that 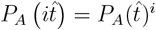. We denote the probability that a mosquito is a HLC in the *i*^*th*^ 15-minute HLC interval, for *i* = 1, 2, …, 18, *p*_*Hi*_. For the first hour of HLCs *p*_*H*1_ = *P*_*H*_ (1*/*4), *p*_*H*2_ = *P*_*A*_(1*/*4)*P*_*H*_ (1*/*4), and *p*_*H*3_ = *P*_*A*_(1*/*4)^2^*P*_*H*_ (1*/*4). For the subsequent hours we assume that HLCs in the first 15-minute interval are a proxy for the HLCs which would have occurred during the last 15-minute interval of the prior hour as well as the first 15-minute interval of the current hour (in which they were caught). Therefore, _*pH*4_ = *P*_*A*_(1*/*4)^3^*P*_*H*_ (2*/*4), *p*_*H*5_ = *P*_*A*_(1*/*4)^5^*P*_*H*_ (1*/*4) and *p*_*H*6_ = *P*_*A*_(1*/*4)^6^*P*_*H*_ (1*/*4), and so on.

The probability that a mosquito is still in state *A* at the end of the 6-hour experiment (6 1-hour time steps) is the probability the mosquito is still host seeking (*p*_*A*_ = *P*_*A*_(1)^6^). However, in the datasets 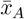 will include disarmed mosquitoes. Therefore, if we consider the the true probability a mosquito remains host seeking is 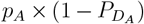. Rearranging for the probability a mosquito is captured resting and survives 24 hours, denoted *p*_*A*_†, gives

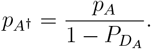

The probability that a mosquito is killed preprandially or is disarmed is *p*_*DM*_ = *P*_*DM*_ (6).

Combining for a vector of the probability a mosquito is a HLC each 15-minute interval, resting, and killed or disarmed during the experiment we get

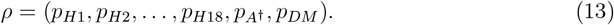

We fit multinomial distributions, with probabilities *ρ*_*k*_[*C*] and *ρ*_*k*_[*I*], to the datasets *D*_*k*_[*C*] and *D*_*k*_[*I*], respectively.

The host-availability rate should only include mosquitoes which are not disarmed. However, some disarmed mosquitoes may have been caught by HLC. We denote the rate of HLCs including disarmed mosquitoes as *α*_*H*_†,*k* [*I*] and the rate of preprandial mortality or disarming excluding disarmed HLC mosquitoes as *α*_*DM*_†,*k* [*I*]. These can then be adjusted for the true host-availability rate and rate of preprandial mortality or disarming given as

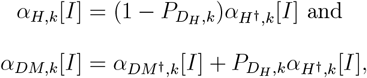

respectively, where 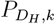 is the nightly probability a HLC mosquito is disarmed.

#### 3.3.3 Postprandial mortality

Since we assume that some HLC mosquitoes may be disarmed, postprandial mortality analysis considers only mosquitoes which feed because these mosquitoes cannot be disarmed (by definition). We assume the postprandial mortality is the same for all HLC mosquitoes which are not disarmed, despite whether they accepted the blood meal.

Given the total number of blood-fed and blood-fed, surviving mosquitoes each experiment are denoted 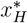 and 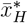, respectively, we denote the nightly values 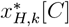 and 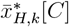 for the control and 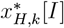and 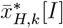 for the intervention experiment, respectively. Then

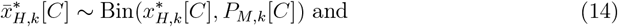

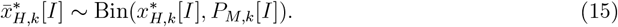

### 3.4 Predicting relative reduction in vectorial capacity

Vectorial capacity is a measure of the ability of the vector population to transmit a disease pathogen, defined as the total number of potentially infectious bites that would eventually arise from all the mosquitoes biting a single perfectly infectious human on a single day [16]. To quantify the effect of an intervention, we calculate the relative reduction of the vectorial capacity when an intervention is utilised in a population compared to when it is not. This is analysed at a range of *coverage levels* within the population. Here coverage refers to the percentage of intervention use within the population. If the intervention is used by 80% of the population, however they only adhere to using it 50% of the time the coverage would be 40%. Here we refer to a host as a host a mosquito may feed upon, rather than a malaria host.

Firstly, the availability rates of unprotected humans and non-human hosts, as well as the per-capita death rates of mosquitoes while searching for a blood meal, are calculated using the model described in Briët et al. [5]. The rates at which mosquitoes encounter human and non-human hosts are scaled to distribute the ratio of bites according to the human blood index (proportion of mosquitoes which feed on humans out of all potential hosts) [5]. Then the relative reduction in vectorial capacity can be calculated using a previously published model [7]. This model assumes the vectorial capacity is proportional to the mosquito emergence rate and therefore is independent of the larval carrying capacity of the environment.

Parameters used in the calculation for the baseline scenario (without the intervention) are given in Table S2. Human hosts in this model are referred to as unprotected hosts. These parameters represent *Plasmodium falciparum* malaria in an *Anopheles minimus* mosquito population. When considering the intervention scenario (with the intervention) we consider an additional human host type with the intervention, protected hosts. Additionally, we include a non-human ’dummy’ host, this facilitates the inclusion of disarming since mosquitoes are assumed to rest for two days after feeding and there is no malaria transmission associated with biting this host. Given the coverage, *C*, the number of protected-human, unprotected-human and dummy hosts are given as *C* × *N*_*H*_, (1 *− C*) × *N*_*H*_ and *C* × *N*_*H*_, respectively. The host-availability rate of protected-human and dummy hosts are calculated as (1 *− π*) × *α*_*human*_ and *κ* × *α*_*human*_, respectively, where *α*_*human*_ is the unprotected-human availability rate. Preprandial killing is included by setting the probability that a mosquito finds a resting place after biting a dummy host to *ω*. Postprandial killing is included by setting the probability that a mosquito finds a resting place after biting a protected host to (1 *− ξ*) × *P*_*C*_. The parameters for the non-human hosts remain the same in the intervention scenarios. To account for the uncertainty in the estimates, we calculate the relative reduction in vectorial capacity for 1000 randomly selected parameter sets from the posterior distribution.

## 4 Results

### 4.1 Intervention parameters

Table 4 gives the median estimates and 95% confidence intervals (CIs) for the relative reduction in the host availability rate (*π*), increase in the rate of disarming or preprandial killing (*κ*), increase in the probability a mosquito is killed postprandially (*ξ*) and proportion of mosquitoes disarmed of all mosquitoes killed predrandially or disarmed (*ω*), for each intervention arm. Generally, the confidence intervals for *κ* are larger than *π*, suggesting there is more uncertainty and/or variability in the change in the preprandial mortality or disarming rate compared to the change in the host-availability rate. Most of the impact of interventions appears to take place preprandially, with relatively low values of *ξ* compared to *π* and *κ*. Many of the larger median values for *ξ* have large 95% CIs, including very small lower bounds. However, the 95% CIs for some interventions with smaller estimated *ξ* also have large upper bounds. Larger CIs may represent more variability, as well as uncertainty.

**Table 4:** Estimated median values and 95% confidence intervals (CIs) of the intervention parameters. *π*: Relative reduction in the host availability rate for mosquitoes encountering a protected human compared to an unprotected human. *κ*: Increase in the rate of preprandial killing or disarming for a protected human compared to the host availability rate of an unprotected human. *ξ*: Increase in the probability of postprandial mortality if a human is protected by the intervention. *ω*: Proportion of mosquitoes disarmed out of all mosquitoes either killed preprandially or disarmed by the intervention.

| Parameter | $\pi$ | $\kappa$ | $\xi$ | $\omega$ |
| --- | --- | --- | --- | --- |
| Round 1 AFRIMS |  |  |  |  |
| EtoCivilianA0 | 0.74 [0.63,0.82] | 0.48 [0.32,0.68] | 0.02 [0.00,0.07] | 0.65 [0.62,0.68] |
| EtoCivilianA20 | 0.53 [0.43,0.61] | 0.24 [0.17,0.31] | 0.01 [0.00,0.02] | 0.91 [0.90,0.92] |
| EtoCivilianAPi0 | 0.87 [0.80,0.92] | 1.65 [1.33,2.04] | 0.11 [0.05,0.20] | 0.05 [0.03,0.07] |
| EtoCivilianAPi20 | 0.73 [0.67,0.78] | 0.44 [0.36,0.53] | 0.02 [0.00,0.06] | 0.63 [0.59,0.65] |
| EtoRanger0 | 0.97 [0.95,0.99] | 0.85 [0.71,1.01] | 0.08 [0.01,0.29] | 0.16 [0.15,0.17] |
| EtoRanger20 | 0.74 [0.65,0.80] | 0.43 [0.32,0.56] | 0.06 [0.01,0.13] | 0.56 [0.51,0.60] |
| EtoRangerPi0 | 0.96 [0.93,0.98] | 0.84 [0.70,1.00] | 0.07 [0.00,0.30] | 0.52 [0.50,0.53] |
| EtoRangerPi20 | 0.82 [0.77,0.86] | 0.57 [0.47,0.68] | 0.01 [0.00,0.05] | 0.73 [0.72,0.75] |
| TransActive | 0.83 [0.77,0.87] | 0.74 [0.59,0.90] | 0.02 [0.00,0.08] | 0.51 [0.47,0.54] |
| TransPassive0 | 0.40 [0.29,0.50] | 0.79 [0.65,0.94] | 0.01 [0.00,0.04] | 0.17 [0.12,0.22] |
| Round 2 AFRIMS |  |  |  |  |
| TransPassive0 | 0.60 [0.54,0.66] | 0.55 [0.47,0.63] | 0.01 [0.00,0.03] | 0.43 [0.36,0.50] |
| MetoPassive0 | 0.42 [0.33,0.50] | 0.51 [0.43,0.60] | 0.12 [0.07,0.17] | 0.39 [0.34,0.44] |
| EtoCivilianBPi0<br>+ TransPassive0 | 0.84 [0.77,0.89] | 2.16 [1.80,2.60] | 0.13 [0.01,0.31] | 0.30 [0.26,0.48] |
| Round 2 KU |  |  |  |  |
| EtoCivilianBPi0 | 0.57 [0.43,0.68] | 1.99 [1.71,2.31] | 0.41 [0.27,0.56] | 0.32 [0.29,0.35] |
| TransPassive30 | 0.98 [0.97,0.99] | 0.42 [0.35,0.50] | 0.11 [0.01,0.34] | 0.78 [0.77,0.81] |
| EtoCivilianBPi0<br>+ TransPassive0 | 0.99 [0.98,1.00] | 0.69 [0.60,0.80] | 0.26 [0.03,0.63] | 0.49 [0.47,0.52] |

Figure 4 compares the effects of each intervention. Figures S1–S3 show distributions of estimated parameters. We observe that washing treated clothes decreases the reduction in biting (*π*), decreases the overall effect on disarming and preprandial mortality (*κ*) and decreases the effect on postprandial mortality (*ξ*). Washing also reduces the contribution of preprandial mortality to *κ*. However, for EtoRanger and EtoCivilianAPi there is an increase in the contribution to *κ* from disarming. For these interventions, as clothes are washed, some mosquitoes that would have been killed by unwashed clothes are disarmed instead. This suggests the response effects to the active ingredients is dose-dependent.

The median predicted probability a HLC mosquito was disarmed 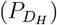 was between 0.12 and 0.91 for intervention-arms. Since the median value for all interventions is larger than 0, if blood feeding was not considered, the reduction in biting would have been underestimated. The median predicted probability that a resting mosquito was disarmed 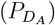 was between 0.04 and 0.57 for intervention-arms. This indicates that disarming in HLCs in more common than resting mosquitoes, however if there are a larger number of resting mosquitoes, the total number of disarmed resting mosquitoes may still be larger. If blood feeding was not considered, these mosquitoes would be considered as repelled and would therefore search for a blood meal elsewhere, which is not the case for disarmed mosquitoes.

### 4.2 Vectorial capacity

The predicted reduction in vectorial capacity for a range of coverage levels is shown in Figure 3. All interventions predict relatively large reductions in vectorial capacity, even at low coverage levels. The estimated reduction in vectorial capacity is similar when considering parameter estimates from different rounds of the study and sites.

**Figure 3.**
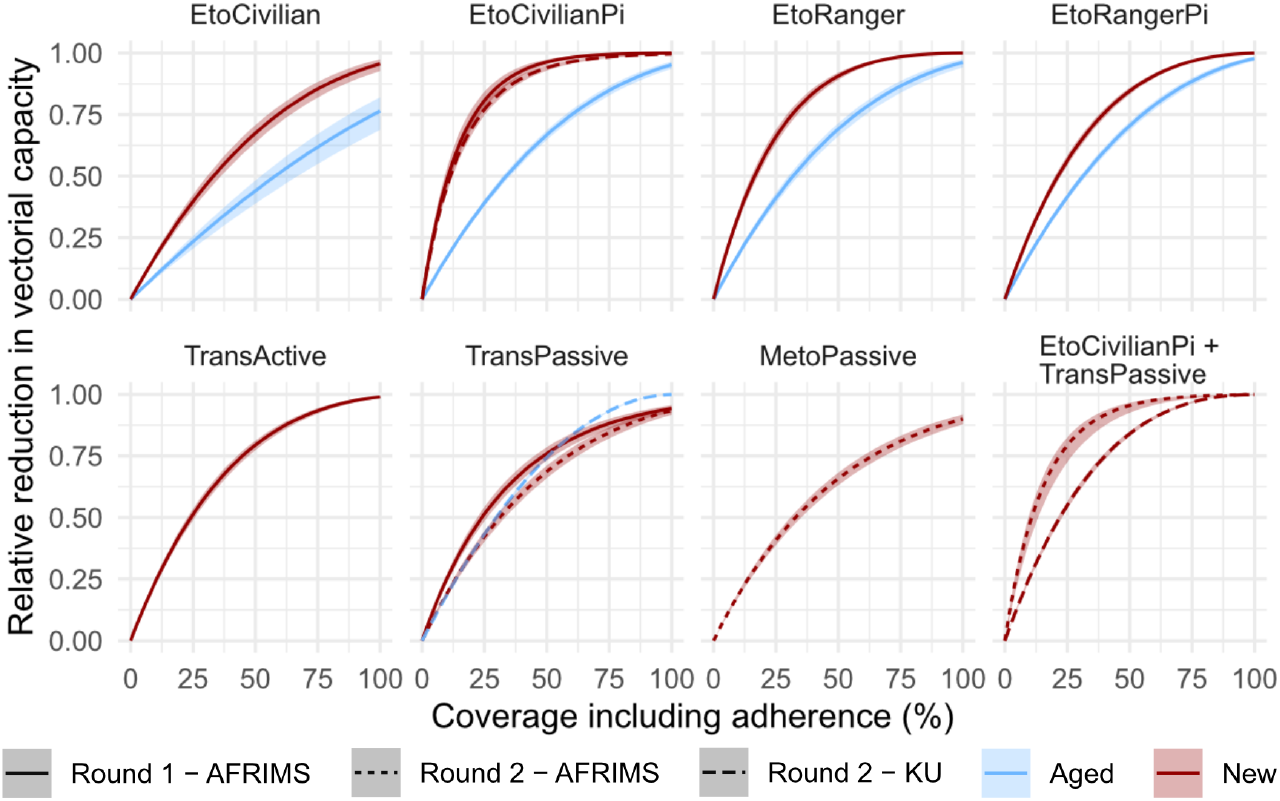
: Relative reduction in vectorial capacity for *Anopheles minimus* due to each intervention for a range of coverage levels. Etofenprox-treated clothing was trailed as new and 20 days old (aged). TransPassive was trialed as new and 30 days old (aged). In round one EtoCivilian type A was used, whereas in round two type B was used.

**Figure 4:**
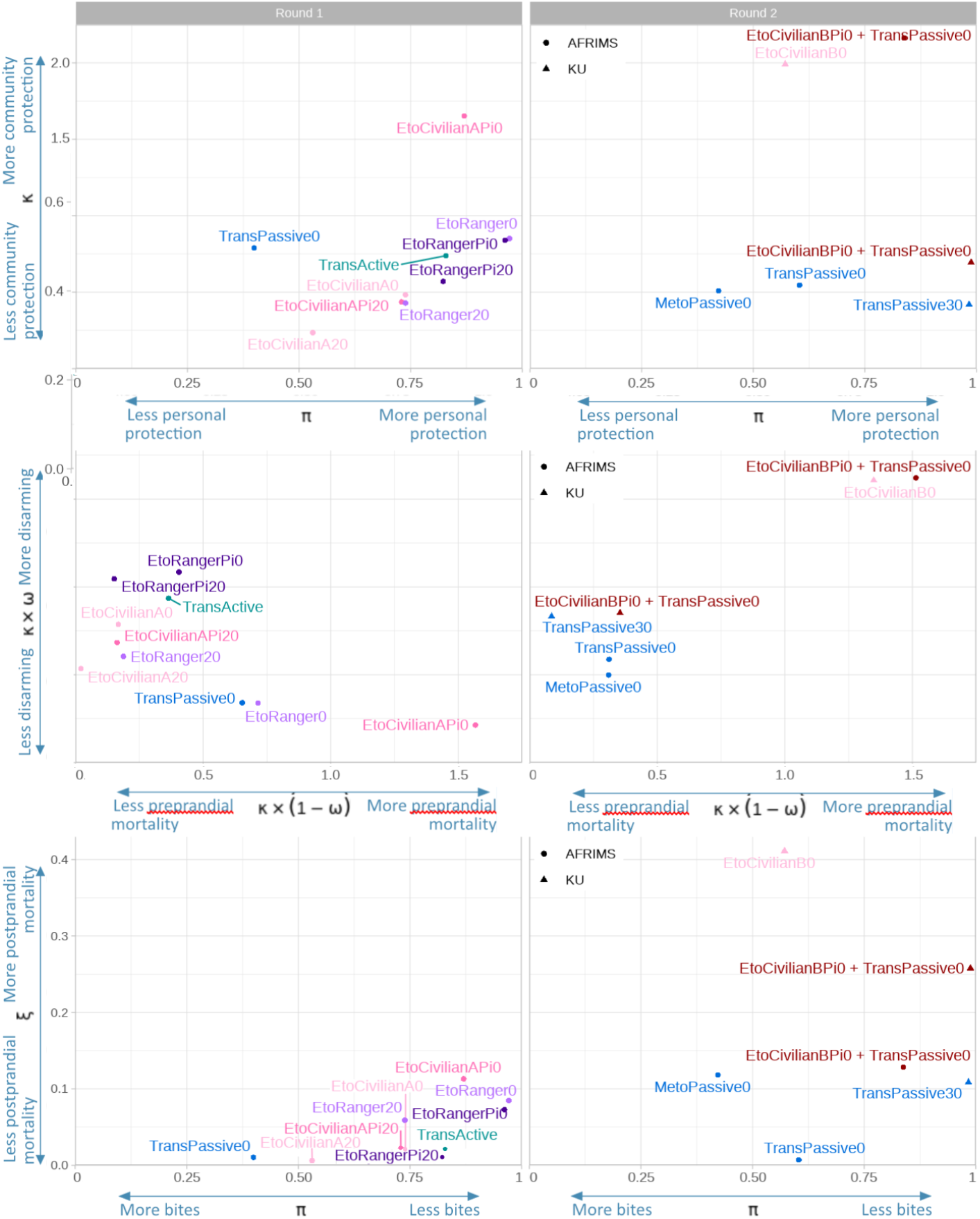
Median estimated values for each intervention of **(Top)** Relative reduction in the host availability rate for mosquitoes encountering a protected human compared to an unprotected human (*π*) and the increase in the rate of preprandial killing or disarming for a protected human compared to the host availability rate of an unprotected human(*κ*), **(Middle)** The contributions to *κ* from disarming (*κ×ω*) and preprandial mortality (*κ×* (1*−ω*)), **(Bottom)** *π* and the increase in postprandial mortality of protected humans compared to unprotected humans (*ξ*).

The etofenprox-treated clothing continues to reduce vectorial capacity when they have been washed 20 times, with relatively small reductions in effectiveness. TransPassive estimates show the aged product as more effective, however new and aged products where not trialled at the same sites.

## 5 Discussion

Most commonly used vector-control interventions, such as LLINs, are treated with pyrethroid insecticides. The physiological resistance to these insecticides in many mosquito vectors, including *Anopheles minimus* [9], is rapidly spreading across malaria endemic countries. New active ingredients that work in the vapour-phase, as well as more rapid and robust approaches to their evaluation, are urgently needed [26]. This analysis shows that etofenprox-treated clothing, with and without picaridin, transfluthrin spatial repellents and metofluthrin spatial repellents performed well in semi-field trials and are promising new vector-control tools. Including mosquito collection by aspiration and allowing for blood-feeding in semifield experiments allowed for the Denz et al. [10] framework to be extended to distinguish between disarming and preprandial killing. Using these parameter estimates we are able to assess both the personal and community protection offered by interventions.

Previously, Syafruddin et al. [33] showed that transfluthrin has potential to reduce disease burden by 66% despite reductions in landing of 16%; the additional community protection characteristics described here could explain this. Similar results have been observed for the impact of transfluthrin on *Aedes*-borne viruses [21].

Although field studies, performed with wild mosquitoes, may provide data on natural behaviour, semi-field studies allow for more detailed analysis of end outcomes of a known mosquito population. The semi-field environment allows for determination of additional modes of action of tools other than reduction in landing or biting. Here we use semi-field experiments to explore the mechanisms behind the reduction in biting, which cannot be observed in a field experiment. While semi-field studies can explain why we see a reduction in biting, this may not be the true reduction of biting that would be observed in a natural setting. However, the further understanding of why the reduction is observed (preprandial killing or disarming) can be combined with field estimates (for the reduction in biting in a natural setting) allowing more accurate estimates of impact of the interventions on vectorial capacity and malaria transmission to consider the community protection of these end points. Analysis showed evidence of disarming in both HLC and resting mosquitoes. Usually, HLCs are assumed to be a proxy for biting, however the reduction in blood feeding is evidence that not all mosquitoes that landed may have bitten [32, 34]. Similarly, if there is a reduction in blood feeding in the resting mosquitoes it may be incorrect to assume these mosquitoes are repelled since some may be disarmed instead. Not including blood feeding would therefore underestimate the reduction in biting for protected hosts and/or underestimate community protections, since it would be assumed that disarmed resting mosquitoes were repelled and would continue to search for a blood meal.

The effectiveness of the killing and disarming effects of some interventions is likely dependent on temperature. Semi-field studies may over-estimate disarming and preprandial mortality since initially repelled mosquitoes remain within a relatively close proximity to the intervention [6]. In the field, these mosquitoes may leave the environment before they become disarmed or dead. Denz et al. [10] suggested that keeping HLC caught mosquito cups in the cage may also have further exposed these mosquitoes to the intervention, therefore in this study the cups were removed every hour.

The model could not be successfully calibrated to all data sets. Round one in AFRIMS had the overall highest proportion of blood fed mosquitoes. For round one, the model could be calibrated to all AFRIMS data sets. However, only EtoCivilianA0 had more than one data point at KU. It is possible that the model calibration failed as one of these days had substantially more human landing catches in the intervention than the control for one of these days [35]. Focusing on round two, the number of data points does not seem to effect whether the model could be calibrated to the data. However, more data points would allow the variability of the effectiveness of the tool to be captured. During round two it was noticed that KU had significantly more knocked-down intervention mosquitoes than AFRIMS. Of these knocked-down mosquitoes more survived at KU, compared to AFRIMS. Figure S4 shows histograms of the proportion recovered and the number of knocked-down mosquitoes that recover after 24 hours at the two sites. This may explain why more disarming was observed at KU. This could either be due to different perceptions of knock down, differences in the interventions (due to human behaviour or temperature), differences in mosquitoes sensitivity to pyrethroids or differences in the fitness of mosquitoes at the two sites.

Though the time-limited (8-day) study demonstrated an increase of TransPassive efficiency over time, the reasons and the duration of this effect are not known. Although the experiments were completed for both new and aged TransPassive in both sites for round two, for AFRIMS and KU only the new and aged products data successfully calibrated the model, respectively. Therefore, it is possible this result is due to a difference between the two sites, for example the mosquito colonies sensitivity or the meteorological conditions. The chemistry of the specific intervention may also be responsible for enhanced volatility at this specific time point, however, more studies are required to understand this phenomenon. This observation points towards a need for product-specific evaluations for understanding how the characteristics of how each product change over time.

The model for vectorial capacity, Chitnis et al. [7], assumes that the intervention does not impact the emergence rate of mosquitoes and that the emergence is only dependent on the larval carrying capacity and that the emergence of mosquitoes is dependent only on this capacity. However, while this might be the case for locations with limited breeding sites compared to the size of the local mosquito population, in locations with high breeding site capacity compared to the size of the mosquito population reducing the number of adult female mosquitoes will also reduce the emergence rate. Therefore, the value represented here is a lower bound for the reduction in vectorial capacity, and in sites with high breeding site capacity the preprandial and postprandial killing effect might further reduce the emergence rate of mosquitoes, and consequently the vectorial capacity [3]. In the calculation of vectorial capacity some of the parameters are derived from another species of *Anopheles* [5], however we do not expect these to be species specific.

Overall, we present a novel statistical framework for estimating the impact of vector-control interventions on the reduction of biting, repellency, preprandial killing, postprandial killing and disarming from entomological semi-field data. Results show how the novel interventions show potential to significantly reduce the vectorial capacity of mosquitoes in the semi-field environment. Extending the standard semi-field trial, from only considering HLC counts, to include aspiration and blood-feeding, increased the observed impact of these interventions for both personal and community protection.

## Acknowledgements

Calculations were performed at sciCORE (http://scicore.unibas.ch/) scientific computing center at University of Basel. The KU Semi field System (SFS) was constructed in the area of Thai Military Development Office, Ministry of Defence and was technically supported by the Kasetsart University Research and Development Institute (KURDI) (Grant No. FF(KU) 14.64).

## Ethical approval

The authors confirm that the ethical policies of the journal, as noted on the journal’s author guidelines page, have been adhered to. No ethical approval was required.

## Author contributions

**Emma L Fairbanks:** Methodology, Software, Validation, Formal analysis, Investigation, Data Curation, Writing -Original Draft, Writing -Review Editing, Visualization, Project administration. **Manop Saeung:** Methodology, Writing -Review Editing. **Arissara Pongsiri:** Methodology, Writing -Review Editing. **Elodie Vajda:** Data Curation, Writing -Review Editing. **Yuqian Wang:** Methodology, Writing -Review Editing. **David J McIver:** Writing -Review Editing, Project administration. **Allison Tatarsky:** Conceptualization, Project administration. **Neil F Lobo:** Conceptualization, Writing -Review Editing. **Sarah J Moore:** Conceptualization, Writing -Review Editing. **Alongkot Ponlawat:** Methodology, Writing -Review Editing. **Theeraphap Chareonviriyaphap:** Methodology, Writing -Review Editing. **Amanda Ross:** Data Curation, Writing -Review Editing. **Nakul Chitnis:** Conceptualization, Resources, Writing -Review Editing, Project administration.

**Financial support** The project was funded by the Malaria Elimination Initiative (A134328). NC and ELF were additionally supported by the Bill and Melinda Gates Foundation (INV025569).

