## Supplementary material for "Inference for entomological semi-field experiments: Fitting a mathematical model assessing personal and community protection of vector-control interventions": Tables

### 1 Supplementary table S1

Table S1: The number of experiments were at least 50% of the control mosquitoes released landed and blood fed when offered a blood meal. Dashes represent interventions were the experiments were not performed or only partly performed (did not include blood feeding). The mean and range of the number of mosquitoes which landed and accepted a blood meal for experiments included in the analysis. † = Model successfully calibrated to the data.

| Intervention | AFRIMS year 1 | KU year 1 | AFRIMS year 2 | KU year 2 |
|---|---|---|---|---|
| EtoCivilianA0 | 2$^{\dagger}$ (76, 54–90) | 2 (54, 51–56) | _ | _ |
| EtoCivilianA20 | 4$^{\dagger}$ (88, 74–96) | _ | _ | _ |
| EtoCivilianAPi0 | 4$^{\dagger}$ (80, 59–93) | _ | _ | _ |
| EtoCivilianAPi20 | 4$^{\dagger}$ (90, 78–100) | 1 (78) | _ | _ |
| EtoCivilianBPi0 | _ | _ | 8 (76, 66–90) | 6$^{\dagger}$ (58, 51–62) |
| EtoRanger0 | 5$^{\dagger}$ (84, 64–97) | _ | _ | _ |
| EtoRanger20 | 3$^{\dagger}$ (80, 53–94) | _ | _ | _ |
| EtoRangerAPi0 | 4$^{\dagger}$ (87, 72–100) | _ | _ | _ |
| EtoRangerAPi20 | 5$^{\dagger}$ (94, 53–78) | _ | _ | _ |
| TransActive | 5$^{\dagger}$ (74, 63–88) | 0 | _ | _ |
| TransPassive0 | 7$^{\dagger}$ (79, 69–90) | 0 | 8$^{\dagger}$ (83, 72–90) | 5 (74, 62–83) |
| TransPassive30 | _ | _ | 8 (89, 67–99) | 7$^{\dagger}$ (67, 59–77) |
| MetoPassive0 | _ | _ | 8$^{\dagger}$ (68,51–81) | 7 (70, 54–82) |
| EtoCivilianBPi0 + TransPassive0 | _ | _ | 8$^{\dagger}$ (79, 64–92) | 7$^{\dagger}$ (68, 51–88) |

### 2 Supplementary table S2

Table S2: Definitions and values of parameters used to calculate the vectorial capacity. Baseline values, considering all humans as unprotected, for *Plasmodium falciparum* malaria and *Anopheles minimus* are given.

| Symbol | Definition | Value | Ref. |
|---|---|---|---|
| $N_H$ | Number of humans | 10,000 | Assumption |
| $N_A$ | Number of non-human hosts | 10,000 | Assumption |
| $\theta_d$ | Maximum time a mosquito unsuccessfully searches for a blood meal per day | 0.33 days | [1] |
| $P_B$ | Probability that a mosquito bites after encountering a host | 0.95 | [1] |
| $P_C$ | Probability that a mosquito finds a resting place | 0.95 | [1] |
| $P_D$ | Probability that a mosquito survives the resting phase | 0.99 | [1] |
| $P_E$ | Probability that a mosquito lays eggs and returns to host-seeking | 0.88 | [1] |
| $\tau$ | Time between feeding and laying eggs | 3 days | [2] |
| $\theta_s$ | Duration of the extrinsic incubation period (time required for sporozoites to develop in the mosquito) | 10 days | [1] |
| $A_0$ | The sac proportion of mosquitoes (estimated proportion of host-seeking mosquitoes which laid eggs in the last 24 hours) | 0.60 | [2] |
| $M$ | The parity proportion of mosquitoes (proportion of host-seeking mosquitoes that have previously laid eggs) | 0.60 | [2] |
| $\chi$ | Human blood index (proportion of blood-fed mosquitoes which fed on a human) | 0.61 | [2] |
| $\alpha_{human}$ | Human availability rate | $4.81 \times 10^{-5}$ days$^{-1}$ | Calculated |
| $\alpha_{non-human}$ | Non-human availability rate | $1.24 \times 10^{-4}$ days$^{-1}$ | Calculated |
| $\mu_{vA}$ | Per-capita mosquito death rate while searching for a blood meal | 0.61 days$^{-1}$ | Calculated |

[2] Y Wang, N Chitnis, and EL Fairbanks. Identification of desirable characteristics of mosquito control tools for reducing malaria transmission in the Greater Mekong Subregion. In preparation.

### 3 Supplementary figure S1

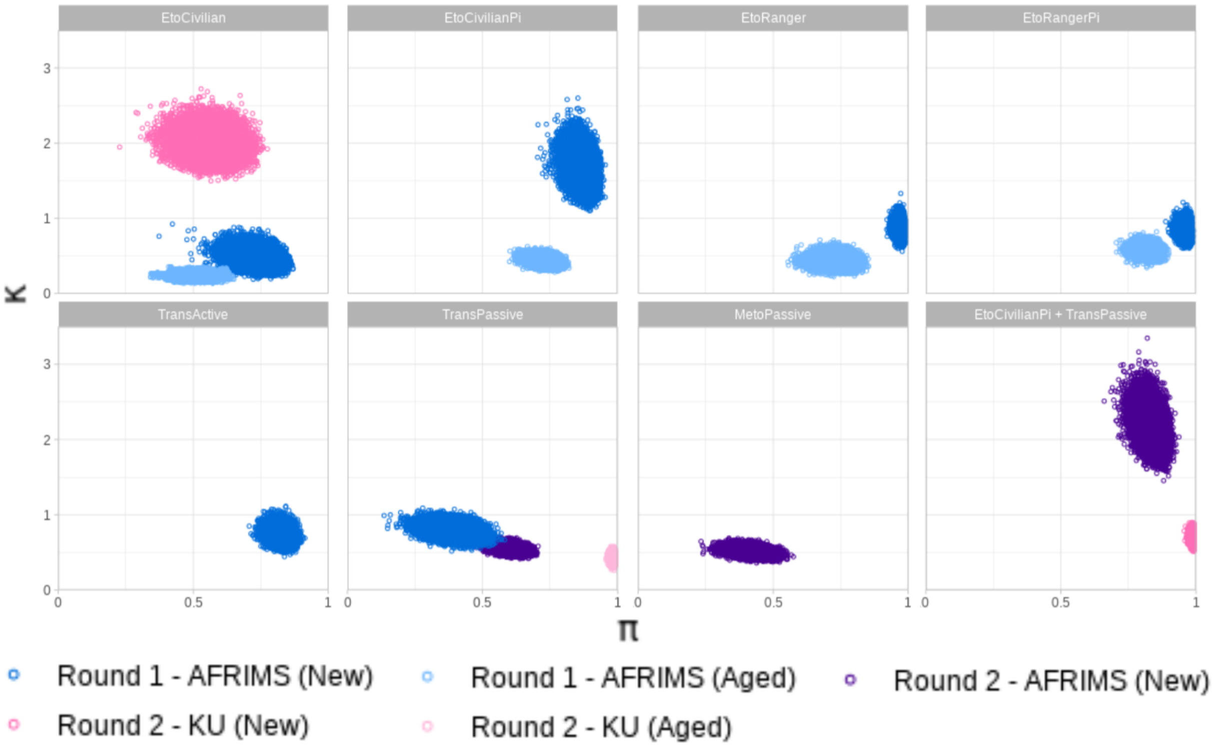


Figure S1: Posterior distributions of $\pi$ and $\kappa$. Etofenprox-treated clothing was trailed as new and 20 days old (aged). TransPassive was trialed as new and 30 days old (aged). In round one EtoCivilian type A was used, whereas in round two type B was used.

### 4 Supplementary figure S2

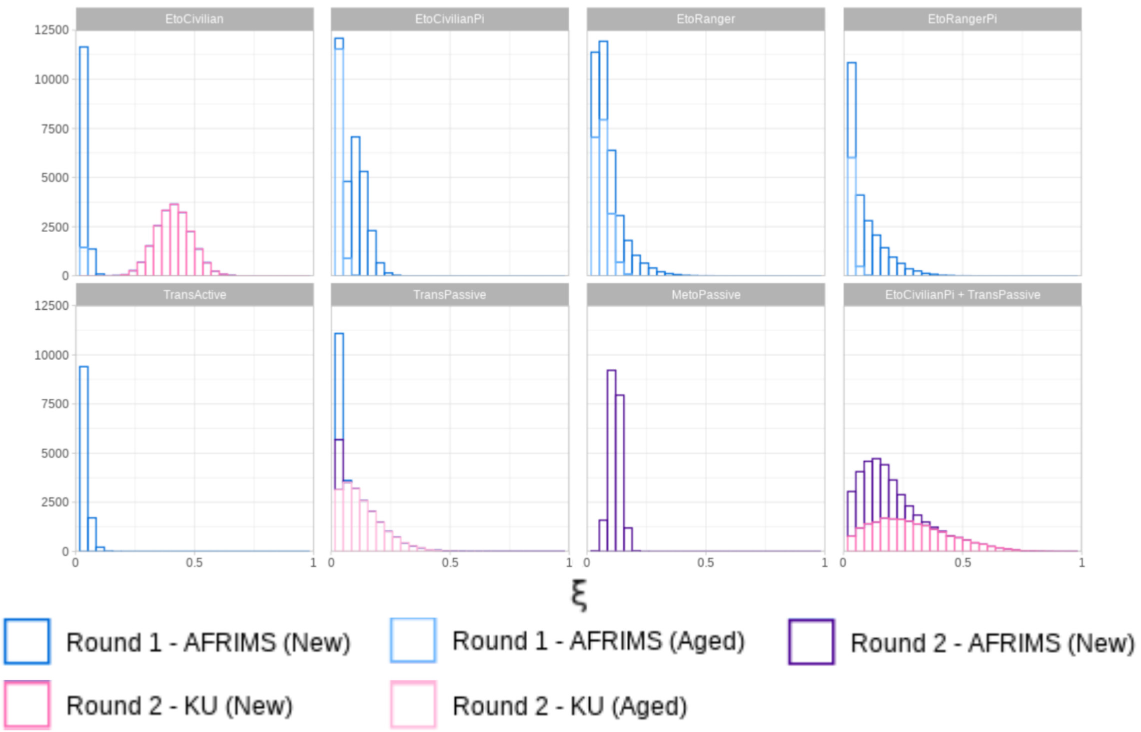


Figure S2: Posterior distribution of $\xi$. Etofenprox-treated clothing was trailed as new and 20 days old (aged). TransPassive was trialed as new and 30 days old (aged). In round one EtoCivilian type A was used, whereas in round two type B was used.

### 5 Supplementary figure S3

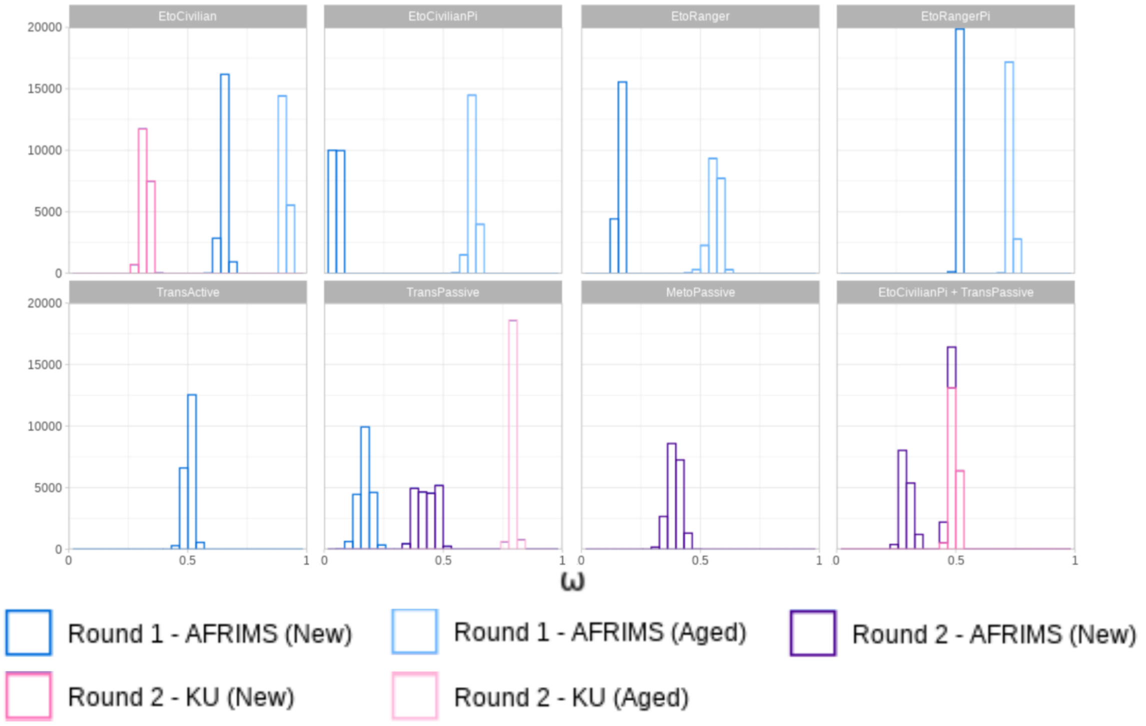


Figure S3: Distribution of estimates of $\omega$. Etofenprox-treated clothing was trailed as new and 20 days old (aged). TransPassive was trialed as new and 30 days old (aged). In round one EtoCivilian type A was used, whereas in round two type B was used.

### 6 Supplementary figure S4

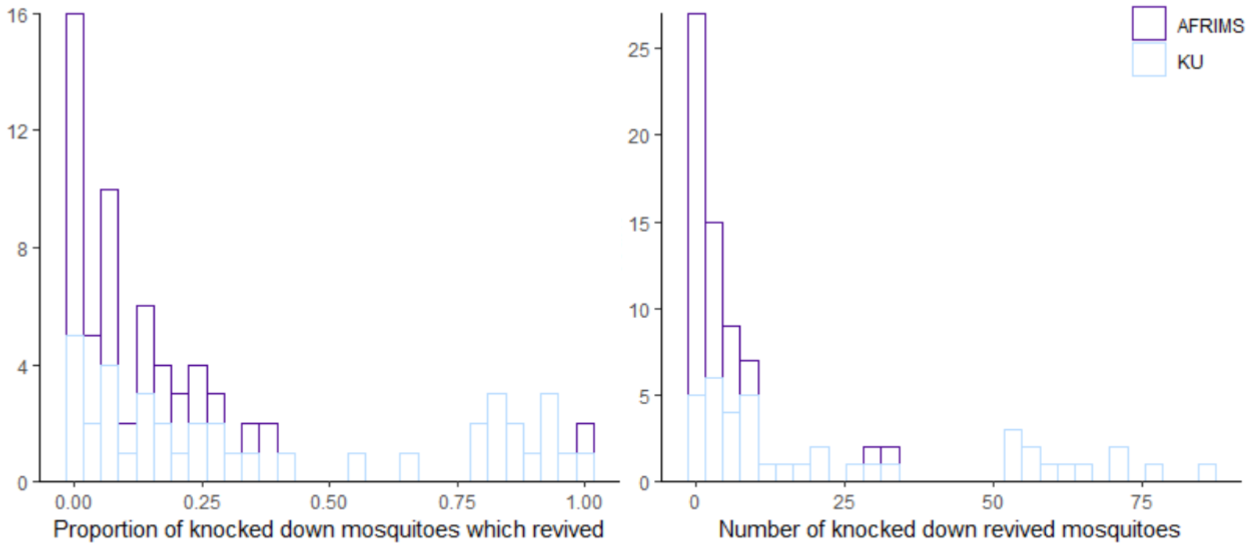


Figure S4: Histograms showing **(Left)** the proportion of mosquitoes knocked down after the six hours of human landing catches which recovered after 24 hours and **(Right)** the number of mosquitoes knocked down after six hours which recovered after 24 hours.
